# A snapshot of SARS-CoV-2 genome availability up to 30^th^ March, 2020 and its implications

**DOI:** 10.1101/2020.04.01.020594

**Authors:** Carla Mavian, Simone Marini, Mattia Prosperi, Marco Salemi

**Affiliations:** Emerging Pathogens Institute, University of Florida, Gainesville, FL, USA; Department of Pathology, University of Florida, Gainesville, FL, USA; Department of Epidemiology, University of Florida, Gainesville, FL, USA

## Abstract

The SARS-CoV-2 pandemic has been growing exponentially, affecting nearly 900 thousand people and causing enormous distress to economies and societies worldwide. A plethora of analyses based on viral sequences has already been published, in scientific journals as well as through non-peer reviewed channels, to investigate SARS-CoV-2 genetic heterogeneity and spatiotemporal dissemination. We examined full genome sequences currently available to assess the presence of sufficient information for reliable phylogenetic and phylogeographic studies in countries with the highest toll of confirmed cases. Although number of-available full-genomes is growing daily, and the full dataset contains sufficient phylogenetic information that would allow reliable inference of phylogenetic relationships, country-specific SARS-CoV-2 datasets still present severe limitations. Studies assessing within country spread or transmission clusters should be considered preliminary at best, or hypothesis generating. Hence the need for continuing concerted efforts to increase number and quality of the sequences required for robust tracing of the epidemic.

**Significance Statement:** Although genome sequences of SARS-CoV-2 are growing daily and contain sufficient phylogenetic information, country-specific data still present severe limitations and should be interpreted with caution.

## Introduction

In December 2019, a novel coronavirus, SARS-CoV-2, was identified in Wuhan, China, as the etiologic agent of coronavirus disease 2019 (COVID-19), which by March 2020 has already spread across more than 80 countries (1). Common symptoms of infection include fever, cough, and shortness of breath, while severe cases are characterized by advanced respiratory distress and pneumonia, often resulting in death (2). It is still unknown what proportion of infected people that only present mild or no symptoms are spreading the virus. A recent study showed that in Wuhan whom roughly 60% of all infections were spread by asymptomatic subjects (3). This characteristic significantly thwarts the job of public health officials who are trying to detect transmission clusters. Few transmission clusters have been identified in China recently through epidemiological contact tracing. However, because of the ongoing nature of the outbreak, it is even more complicated if not impossible to detect transmission clusters using genetic data.

Soon after the first epidemiological and SARS-CoV-2 genetic sequence data were made available, a glut of phylogeny-based analyses began to circulate discussing, in scientific papers as well as (social) media, countries that might have been fueling the spread. The implications of misunderstanding the real dynamic of the COVID-19 pandemic are extremely dangerous. Ethnic or social discrimination resulting from unsupported assumptions on viral contagion – often amplified by irresponsible, uncontrollable communications – can be highly damaging for people and countries. In particular, the US-based NextStrain (4) team has been posting real-time updates on the epidemic tracing by molecular analyses. Despite (social) media are often vehicle for fake news and boast news hype, it is also worth noting the tremendous effort of the scientific community to provide free, up-to-date information on ongoing studies, as well as critical evaluations. Several discussions and evidence-based debates on controversial hypotheses on the epidemic have ensued — such as the number of untraced infections in the US, the putative virus introduction in Italy through Germany, and the alleged lineage diversification in China (5) later criticized (6). Recently, an editorial published on Science (7) has also highlighted how unsupported or misleading claims circulating in forums, social media, and even peer-reviewed articles, have been led by a substantial over interpretation of the available data. Hence, the urgency to reframe the current debate in more rigorous scientific terms, and quantitatively evaluate whether sufficient information for reliable phylogenetic and phylogeographic studies currently exists, or which gaps need to be addressed. We explored the characteristics of few datasets through time and assess phylogenetic signal to understand whether we this data is useful of not.

## Results

### Sampling and phylogeographic uncertainly

Before carrying out any phylogeny-based analysis of virus evolution and spatiotemporal spread, it is crucial to test the quality of sequence data, since uneven sampling, presence of phylogenetic noise, and absence of temporal signal can affect reliability of the results (e.g. ancestral state reconstructions, molecular clock calibrations) (8). SARS-CoV-2 full genome sequences were obatained from GISAID (https://www.gisaid.org/) (9) at different timepoints. As of March 30^th^, we compared the number of full genomes sampled per country with the number of confirmed cases at the time of sampling, as well as the country’s total population (Figure 1). We obtained 2608 full genomes from 55 countries (Figure 1). During the past month, the number of genomes has steeply been increasing: March 3^rd^, 169 genome sequences from 22 countries; March 10^th^, 331 genome sequences from 29 countries; March 18^th^, 794 genome sequences from 35 countries, ρ=0.45; March 25^th^, 1662 genome sequences from 42 countries. We found spearman (rank) correlation between confirmed cases and genomes per country to be 0.49 on March 30^th^, and we considered it as a proxy for sampling homogeneity. However, correlation could only be investigated with confirmed cases (again as proxy), since not all affected countries have made publicly available the total number of coronavirus testing performed. Moreover, even within the same country, sequenced genomes were usually sampled from few hotspots, not necessarily representative of the whole epidemic in that country. It is worrisome that, as of March 30^th^ 2020, the two top countries in terms of confirmed cases do not show sufficiently large and representative sampling. SARS-CoV-2 full genome sequences available from patients in the US, the country with the highest number of confirmed cases, have mainly been sampled in Washington state (66%) during the early epidemic, while less than one third (32%) are available from the epicenter of the US epidemic, the state of New York. Italy, the second country per confirmed cases, uploaded 26 genomes, of which one from the Marche region, four from Friuli Venezia Giulia, seven from Abruzzo, nine from Lazio, and only five from Lombardy, which is epicenter of the Italian epidemic (Table S1) (10). The top 10 contributors per number of genomes are USA (612), Iceland (343), UK (321), China (300), Netherlands (190), France (119), Japan (83), Canada (80), Australia (64), and Belgium (46). Notably, some countries uploaded a high number of genomes despite having a relatively low number of cases (e.g., Georgia, Iceland, Senegal, DRC).

**Figure 1.**
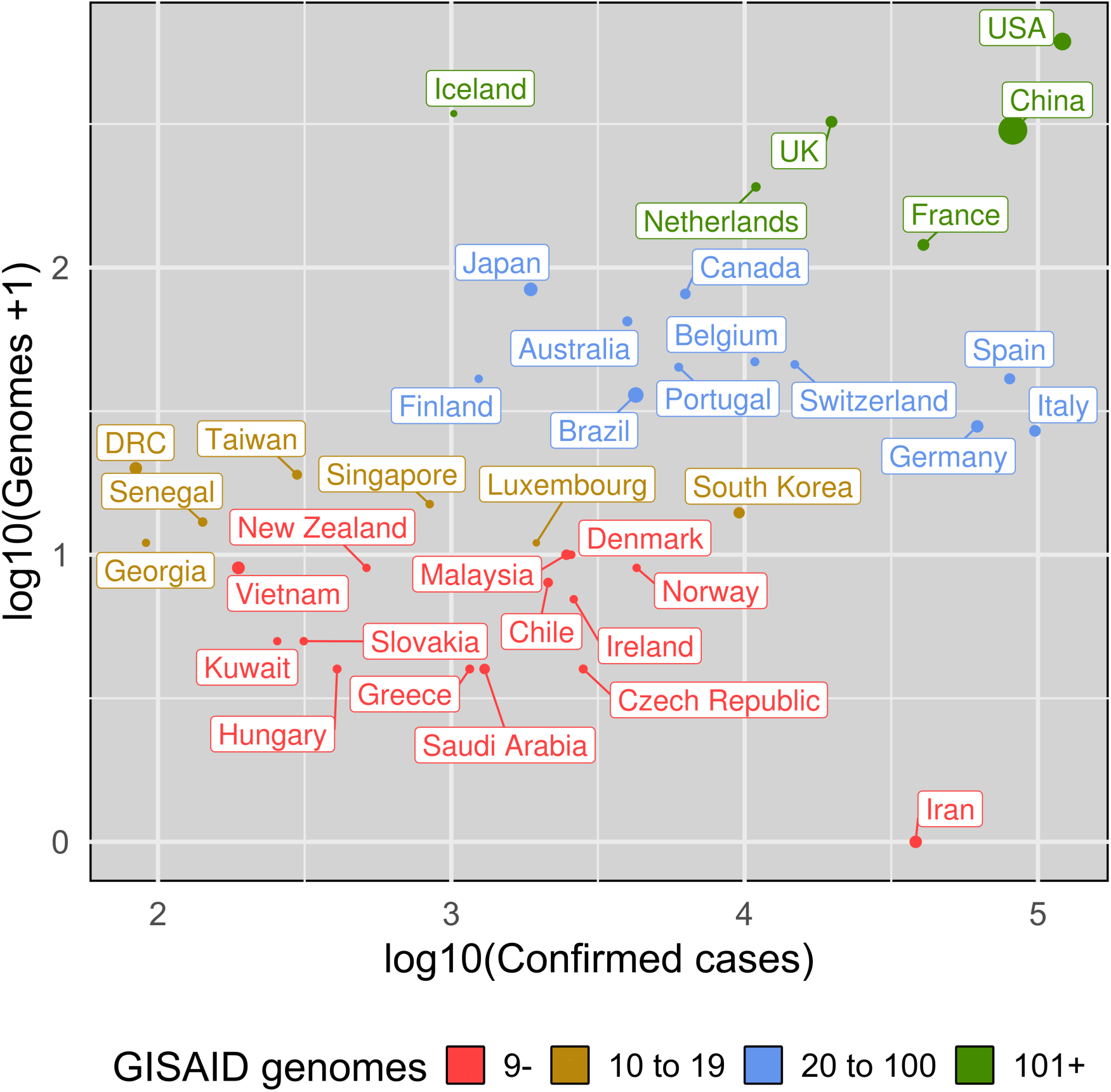
Available SARS-CoV-2 genome sequences and confirmed cases per country. Selected countries for number of provided GISAID full genomes (2 or more), or confirmed cases (25K or more) as of March 30th 2020. On the x axis we have number the confirmed cases (log10 scale), on the y axis the number of GISAID full genomes +1 (log10 scale). The number of genomes determines dot color, while the country population determines dot size. Notably, some countries uploaded a high number of genomes despite having a relatively low number of cases (e.g., Georgia, Iceland, Senegal, DRC).

### Phylogenetic noise in sequence data

Lack of resolution and uncertainty in the SARS-CoV-2 phylogenetic tree is to be expected, considering that relatively little genetic diversity can be accumulated during the first three months of an epidemic, even for an exponentially spreading and fast-evolving RNA virus. Assessment of phylogenetic signal in the dataset was carried out using likelihood mapping analysis (11), which estimates the likelihood of each of possible tree topology for any group of four sequences (quartet), randomly chosen from an alignment, and reports them inside an equilateral triangle (the likelihood map) where the corners represent distinct tree topologies and the center star-like trees. Quartets are considered “resolved” when the three likelihoods are significantly different (phylogenetic signal, most dots equally distributed in the corners, i.e. data are suitable for robust phylogeny inference), unresolved or partially resolved, when all three likelihood values or two of them are not significantly different (phylogenetic noise, most dots in sides or center areas, i.e. data may not be sufficient for robust phylogeny inference). Extensive simulation studies have shown that, for sequences to be considered robust in terms of phylogenetic signal, side/center areas of the likelihood mapping must include <40% of the unresolved quartets (12). Overall, phylogenetic signal in the present data has been increasing with number of genomes been released. Percentage of unresolved quartets detected in the SARS-CoV-2 full genomes alignment on March 3^rd^ and 10^th^ was still too high to allow reliable inferences (Figure S1). In other words, such a lack of phylogenetic signal has likely resulted in overall unreliable topology of any SARS-CoV-2 tree obtained using those data, and even clades with high bootstrap values should have been interpreted with extreme caution.

The effect of inhomogeneous sampling, lack of phylogenetic signal and missing data on phylogeography reconstructions, like the ones recently rushed through news and (social) media to claim specific dissemination routes of SARS-CoV-2 among countries, can be quite dramatic. An instructive example is the putative introduction of SARS-CoV-2 in Italy from Germany. A preliminary maximum likelihood (ML) tree, inferred from the full genome viral sequences available on March 3^rd^ 2020, showed a well-supported cluster of European and Asian sequences (reported in Figure S2), which contained a subclade (Subclade A, Figure 2a) including a sequence isolated in Germany that appears to be paraphyletic (with strong bootstrap support) to an Italian sequence clustering, in turn, with sequences from Finland, Mexico, Germany and Switzerland. Based on this observation (which was available on NextStrain), a heated discussion circulated on social media about a transmission event from Germany to Italy followed by further spread from Italy the other countries. However, in a new tree inferred just one week later, when more than 135 new full genome sequences were made available on GISAID (9), the direct link between Germany and Italy in Subclade A disappeared due to the additional clustering of previously unsampled sequences from Portugal, Brazil, Wales and Netherland (Figure 2b). In addition, likelihoods of alternative tree topologies generated arbitrarily switching branches in the tree (arrows in Figure 2b), implying different dissemination scenarios, were not significantly different (Shimodaira-Hasgawa test, Table S2) than the likelihood of the tree inferred from the real data. In other words, it is not possible, with present data, to decide which branching pattern (and, therefore, phylogeographic reconstruction) is the one most likely representing actual dissemination routes among European countries.

**Figure 2.**
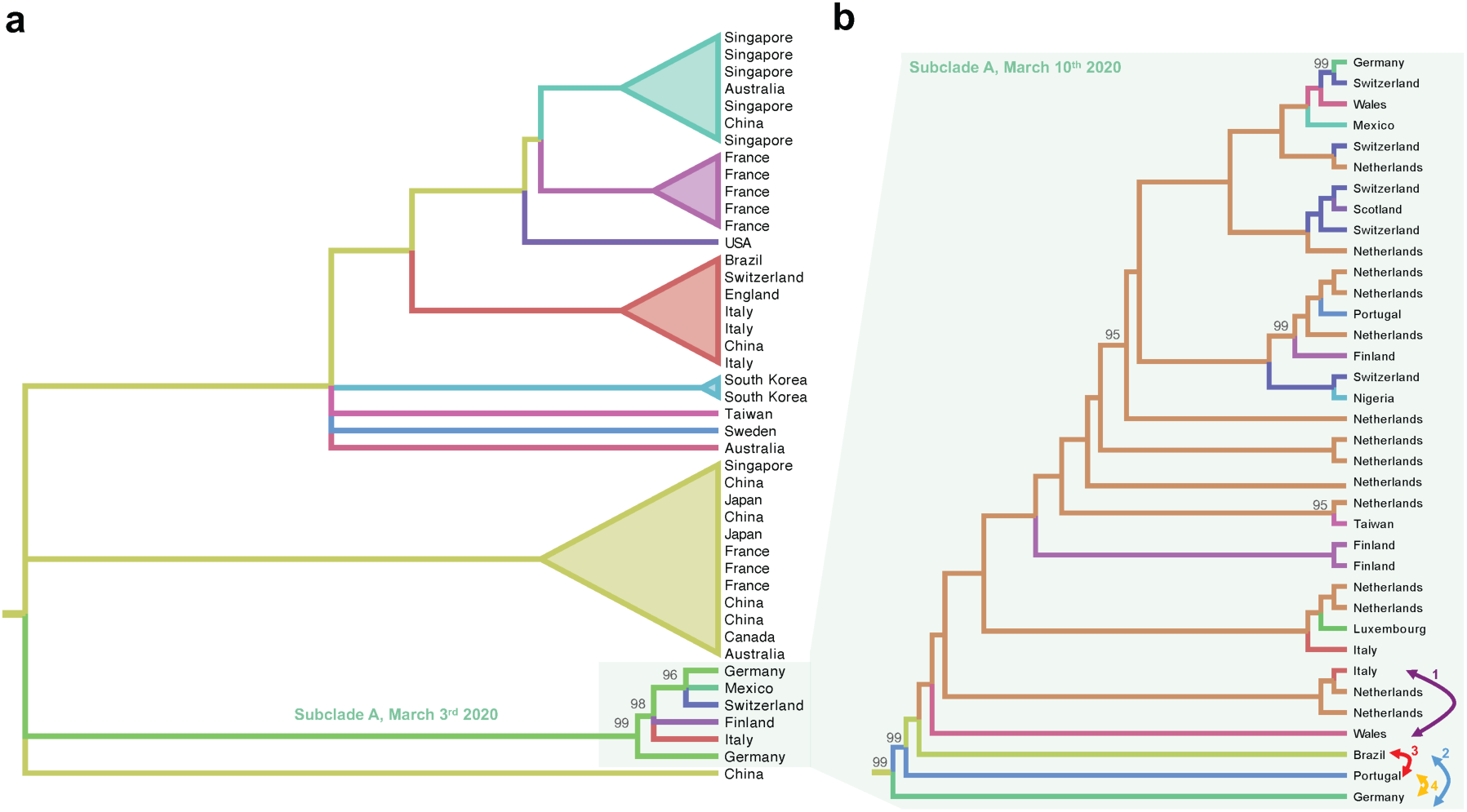
Cladograms of SARS-CoV-2 subclades. Cladograms were extracted from ML phylogenies rooted by enforcing a molecular clock. Colored branches represent country of origin of sampled sequences (tip branches) and ancestral lineages (internal branches). Numbers at nodes indicate ultrafast bootstrap (BB) support (only >90% values are shown). (a) Cladogram of a monophyletic clade within the SARS-CoV-2 ML tree inferred from sequences available on March 3^rd^ 2020 (Supplementary Figure S1). The subclade including sequences from Italy and Germany, named Subclade A, is highlighted. (b) Cladogram of subclade A of the SARS-CoV-2 ML tree including additional sequences available on March 10^th^ 2020 (Figure S2). Each bi-directional arrow, and corresponding number, connects two tip branches that were switched to generate an alternative tree topology to be tested (see Table 1, Methods).

Available genome sequences are rapidly growing. SARS-CoV-2 full genome dataset is now showing less than 40% of unresolved quartets in the center: 38.6% unresolved quartets on March 18^th^ (794 genome sequences) (Figure S1c) and 32.3% on March 25^th^ (1,660 genome sequences) (Figure S1d). This indicates that the amount of phylogenetic information is now potentially usable to define phylogenetic relationships among strains. Plotting mean genetic distance of each sequence from the root of a phylogeny versus the sequence sampling time allows to testing for significant linear correlation, which is necessary for the calibration of a reliable molecular clock (13) (Figure S3). As expected in genomes obtained over a very short period of time (∼ three months) since the beginning of the outbreak, correlation in the current data is fairly week (Table S3). However, Bayesian analysis (14), which infers phylogenetic and phylogeographic patterns from a posterior distribution of trees, might facilitate comparisons about different evolutionary scenarios, help in retrieving the correct topology, and estimate an accurate evolutionary rate using relaxed clock methods (15). Reconstructing the phylogenetic relationships of the same European Subclade A discussed above with sequence available on March 18^th^ 2020 showed a much more complex snapshot of SARS-CoV-2 spreading (Figure S4). Taking a closer look at the Sublade A reveals that even with more genomes available, inference is bias by over-sampling of some countries and under-sampling of others (Figure S4). Yet, even with more genomes available, inference is bias by over-sampling of some countries and under-sampling of others (Figure S4). Recently, methods were developed to estimate, for each pair of viral sequences from two infected individuals, how many intermediates there are in the putative transmission chain connecting them, using a transmission matrix (16). The analysis of SARS-CoV-2 genomes shows that numerous links among samples are still missing (Figure S5). In such a scenario, it is not advisable to extrapolate conclusions on the origin and dissemination of strains.

Phylogenetic signal is increasing in the global alignment; yet, likelihood mapping per country, using data from countries reporting highest number of cases – USA, Italy, Spain, Germany, and France – indicate that some local datasets lack sufficient signal (Figure S6). Lack of signal was found in datasets from Italy (26 genomes, 45 variant sites – 0.2% of total sites in the genome - 11 parsimony informative), USA (612 genomes, 675 variant sites - 2.3% of total sites in the genome - 158 parsimony informative) and China (300 genomes, 742 variant sites – 2.5% of total sites in the genome - 98 parsimony informative). The top 5 contributing states in US are Washington (405, 66%), California (45, 7%), Minnesota (33, 5%), Wisconsin (29, 5%), and Utah (22, 4%); 42 genomes are not labeled with a state or city. USA dataset comprised mostly of sequences collected in Washington State (423 genomes, 69.1%). The top 5 contributing provinces in China are Shanghai (96, 32%), Guangdong (80, 27%), Hong Kong (30, 10%), Hubei (31, 10%), Hangzhou (9, 3%), and Shandong (9, 3%); 20 genomes are not labeled with a province or city. Neither China nor US showed phylogenetic signal despite the high number of genome sequences available (Figure S6). On the contrary, and unexpectedly, countries with low number of genome sequences did show presence of phylogenetic signal: Germany, Spain and France (Figure S6).

Despite the presence of phylogenetic signal in these countries, only genomes form France also show temporal signal that would allow for calibration of a molecular clock and re-framing phylogenetic and phylogeography inferences in spatiotemporal dimension. On the other hand, the transmission matrix for France indicates that considerable links are still missing due to unsampled infected individuals, limiting the reliability of transmission cluster studies based on sequence data (Figure S8).

### Conclusions and Future directions

As more genome sequences, sampled at different time points and from diverse geographic areas, are daily becoming available, in depth Bayesian phylodynamic and phylogeography analyses of the COVID-19 pandemic may soon be a viable option. As long new data do not increase phylogenetic and temporal signal, results will remain highly questionable. Characterization of transmission events is fundamental to understand the dynamics of any infectious disease. From a public health standpoint, being able to trace transmission at the local level is crucial. Within country identification of active transmission clusters would open the way to more effective public health measures. The most optimal inference of transmission events would have a combination of genetic and epidemiological data for a joint analysis. However, it is not possible, at the moment, to identify transmission clusters within regions, counties, or cities, solely on genetic data, since micro-scale genetic data is not yet available. Indeed, transmission investigations that have been performed so far have been based on contact-tracing, epidemiological and clinical data (17, 18).

Published scientific data and media are, nowadays, easily accessible to a worldwide audience; properly weighing the information being shared is important more than ever. We deem that current molecular epidemiology data are not solid enough to provide a scientifically sound analysis of SARS-CoV-2 spread. Despite overall increasing of phylogenetic and temporal signal, we suggest that any conclusion drawn, at present, about existing lineages and direction of viral spread, based on phylogenetic analysis of SARS-CoV-2 sequence data, should be considered at the very best preliminary, and hypothesis-generating. The evolutionary dynamics of SARS-CoV-2 spread is unveiling an unprecedented amount of information, essential to make policy decisions. The whole of humanity is threatened by the current pandemic, and policy makers need to adjust their mitigation measures while the pandemic itself is developing. Some of the urgent answers required lie in the timely availability of abundant, high quality genetic data not only from countries experiencing a high number of reported cases, but also from others that seem to be experiencing, at least for now, a lower number of infections.

## Methods

### Data

GISAID was accessed on March 30^th^ 2020 (Table S1). After quality control of sequences that were not full genomes or contained extensive stretches of unknown nucleotides, March 30^th^ March 30^th^ 2020 of 2608 from 55 countries. Confirmed cases were retrieved from the March 29^th^ data of the Covid19 website provided by Johns Hopkins University (16). Top ten countries per confirmed cases were USA (121,146), Italy (97,689), China (82,122), Spain (80,110), Germany (62,095), France (40,708), Iran (38,309), UK (19,778), Switzerland (14,829), and Netherlands (10,930).

### Phylogenetic signal and ML phylogeny inference

Transition/transversions vs. genetic distance plot were calculated using DAMBE6 (19). Evaluation of the presence of phylogenetic signal satisfying resolved phylogenetic relationships among sequences was carried out with IQ-TREE, allowing the software to search for all possible quartets using the best-fitting nucleotide substitution model (11). ML tree reconstruction was performed in IQ-TREE based on the best-fit model chosen according to Bayesian Information Criterion (BIC) (20, 21). Exploration of temporal structure, i.e. presence of molecular clock in the data, was assessed by regression of divergence -root-to-tip genetic distance-against sampling time using TempEst (13). In this case, absence of a linear trend indicates that the data does not contain temporal signal and that the data is not appropriate for phylogenetic inference using molecular clock models. TransPhylo R package was used to infer transmission matrices of SARS-CoV-2 (16).

## Acknowledgments

MS is supported in part by the Stephany W. Holloway University Chair in AIDS Research. We thank all those who have contributed SARS-CoV-2 genome sequences to the GISAID database (https://www.gisaid.org/).

## Supplementary Figures and Tables

**Figure S1.**
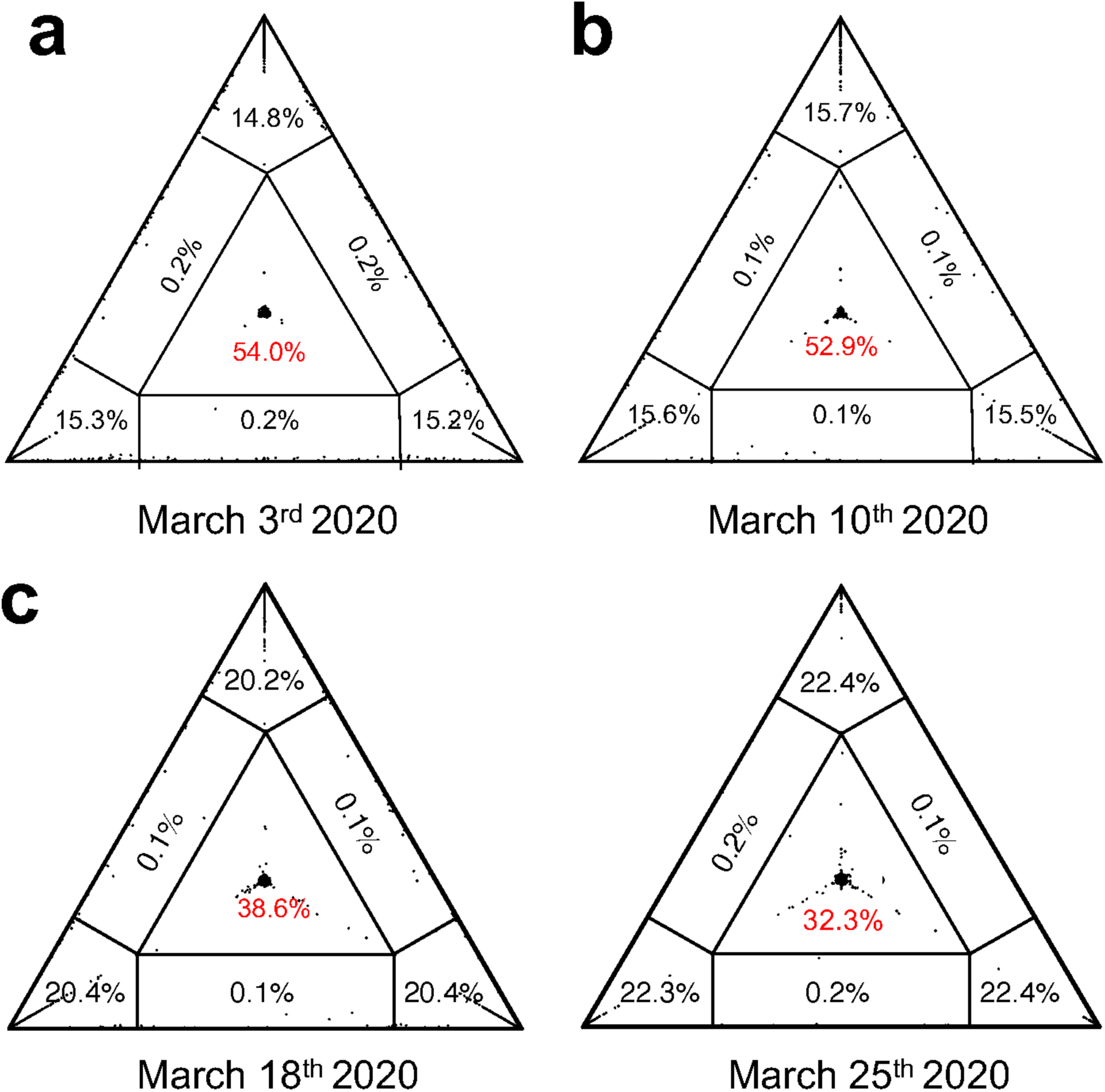
Phylogenetic signal for 169 and 331 full genomes of SARS-CoV-2. Presence of phylogenetic signal was evaluated by likelihood mapping checking for alternative topologies (tips), unresolved quartets (center) and partly resolved quartets (edges) for the (a) 169 genomes available on March 3^rd^ 2020, (b) and 331 genomes on March 10^th^ 2020, (c) 794 genome on March 18^th^ 2020, and (d) 1,660 genome on March 25^th^ 2020.

**Figure S2.**
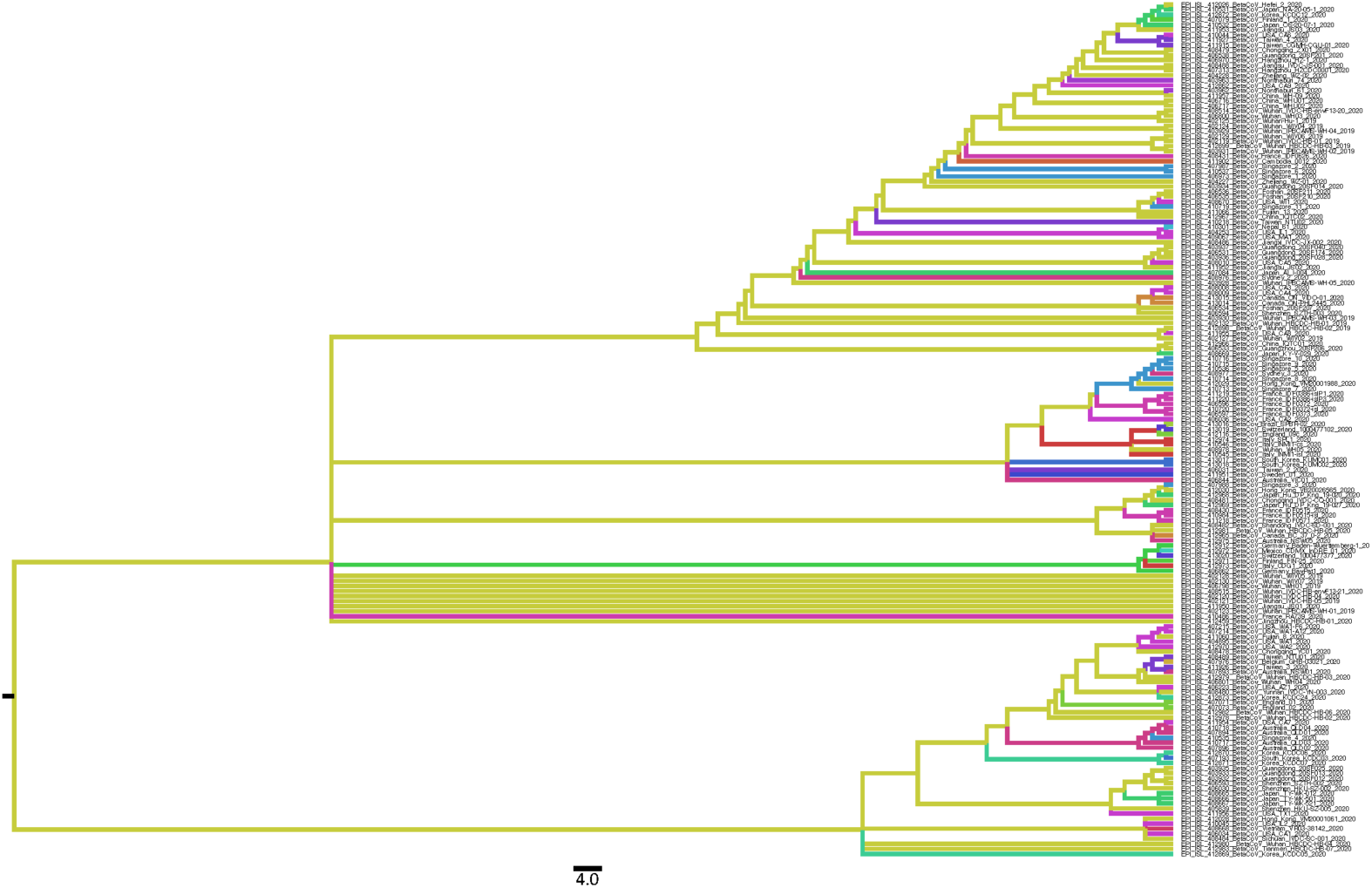
Phylogeographic cladogram of SARS-CoV-2 on March 3^rd^ 2020. Ancestral state reconstruction performed with TreeTime on 169 full genomes of SARS-CoV-2 collected on March 3^rd^ 2020. Branches are colored by countries of origin.

**Figure S3.**
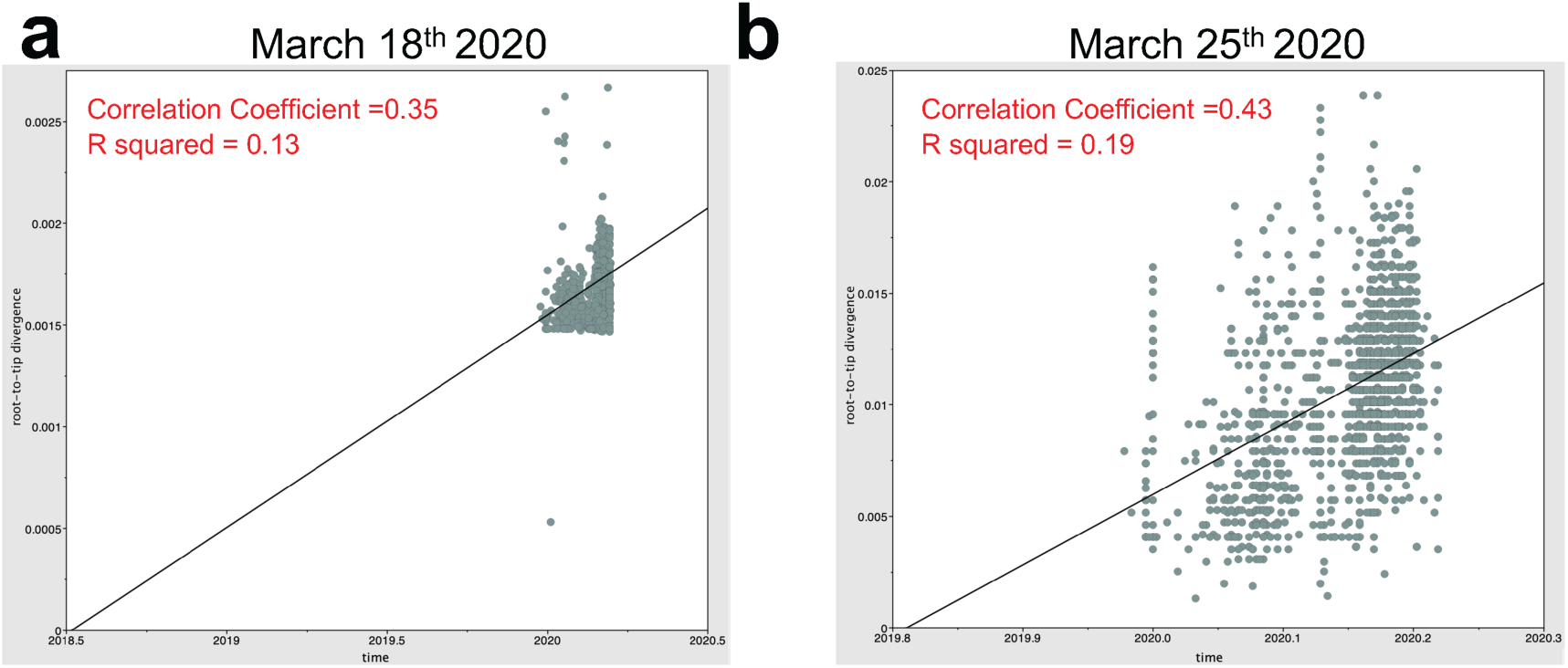
Regression analysis of temporal resolution of SARS-CoV-2 genomes on March 18^th^ and March 25^th^ 2020 datasets. The plots represent linear regression of root-to-tip genetic distance within the ML phylogeny against sampling time for each taxa. Temporal resolution was assessed using the slope of the regression, with positive slope indicating sufficient temporal signal for datasets collecet on (a) March 18^th^ and (b) March 25^th^ 2020. Correlation coefficient “r” are reported for each genomic fragment.

**Figure S4.**
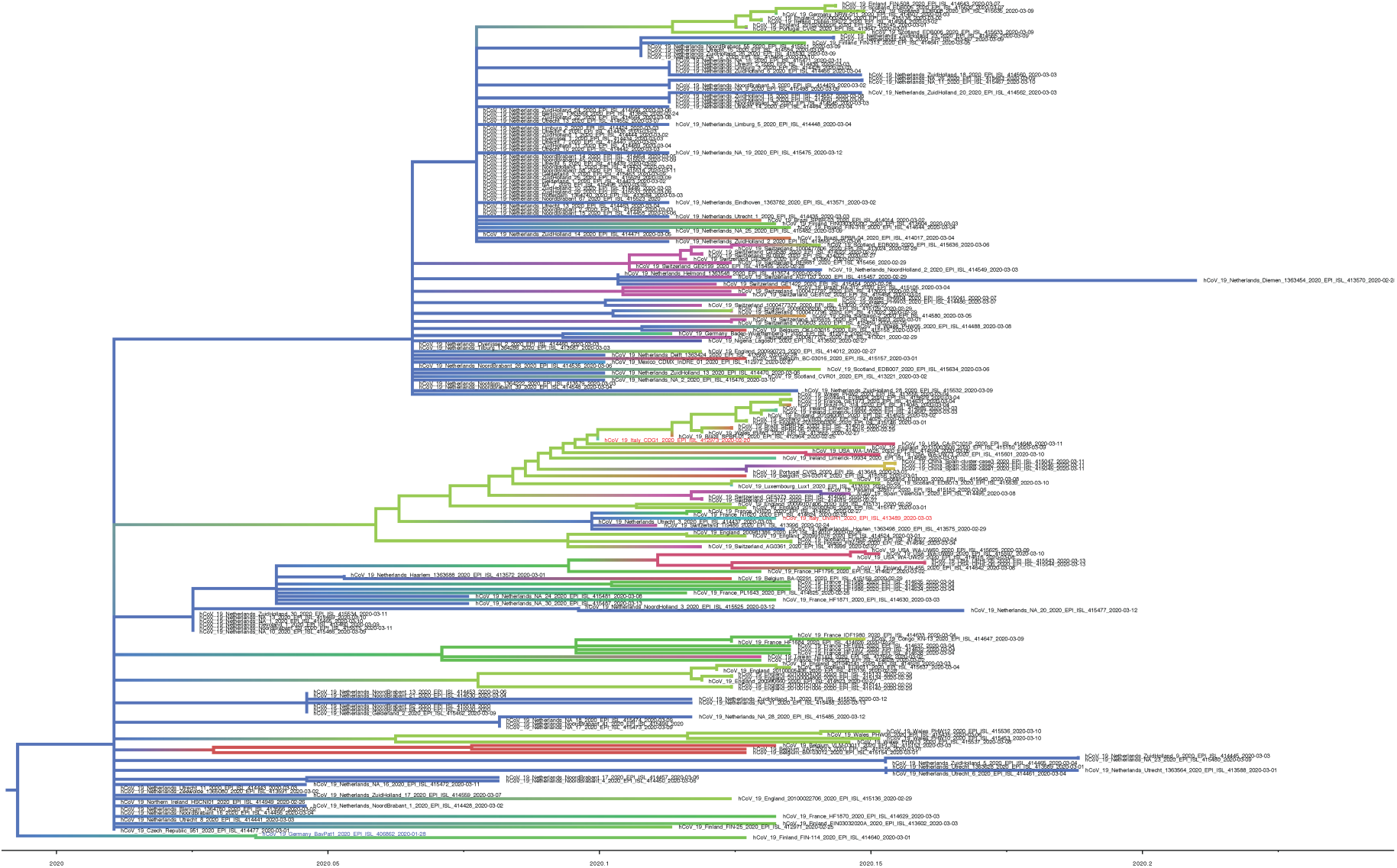
Phylogeographic reconstruction of Subclade A clade as of March 18th. Ancestral state reconstruction performed with TreeTime on Subclade A with genome collected on March 18^th^ 2020. Branches are colored by countries of origin.

**Figure S5.**
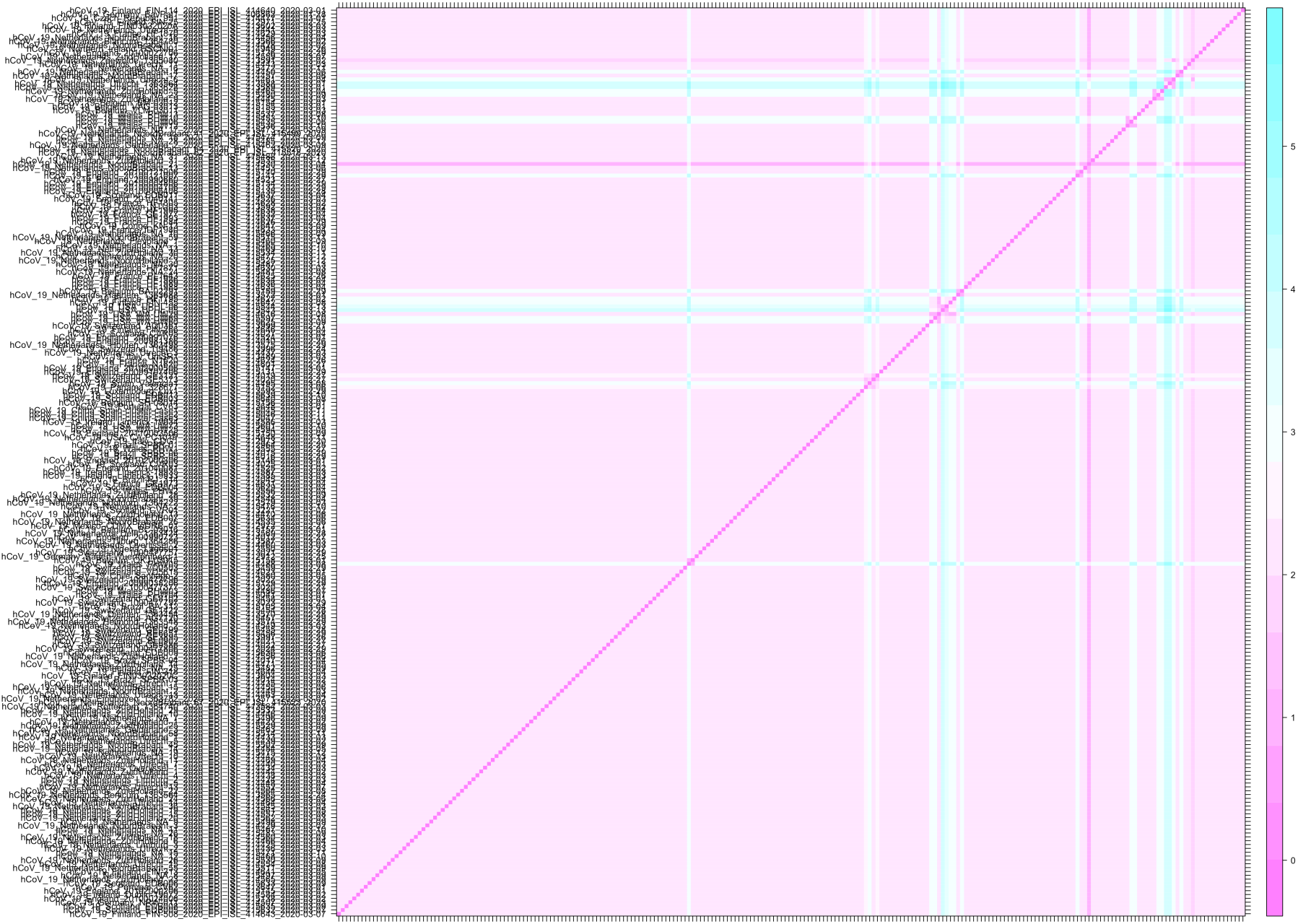
Transmission matrix of SARS-CoV-2 of Subclade A as of March 18^th^ 2020. Transmission matrix indicating for each pairs of individuals how many intermediates there are in the transmission chain are given for Subclade A as of March 18^th^ 2020.

**Figure S6.**
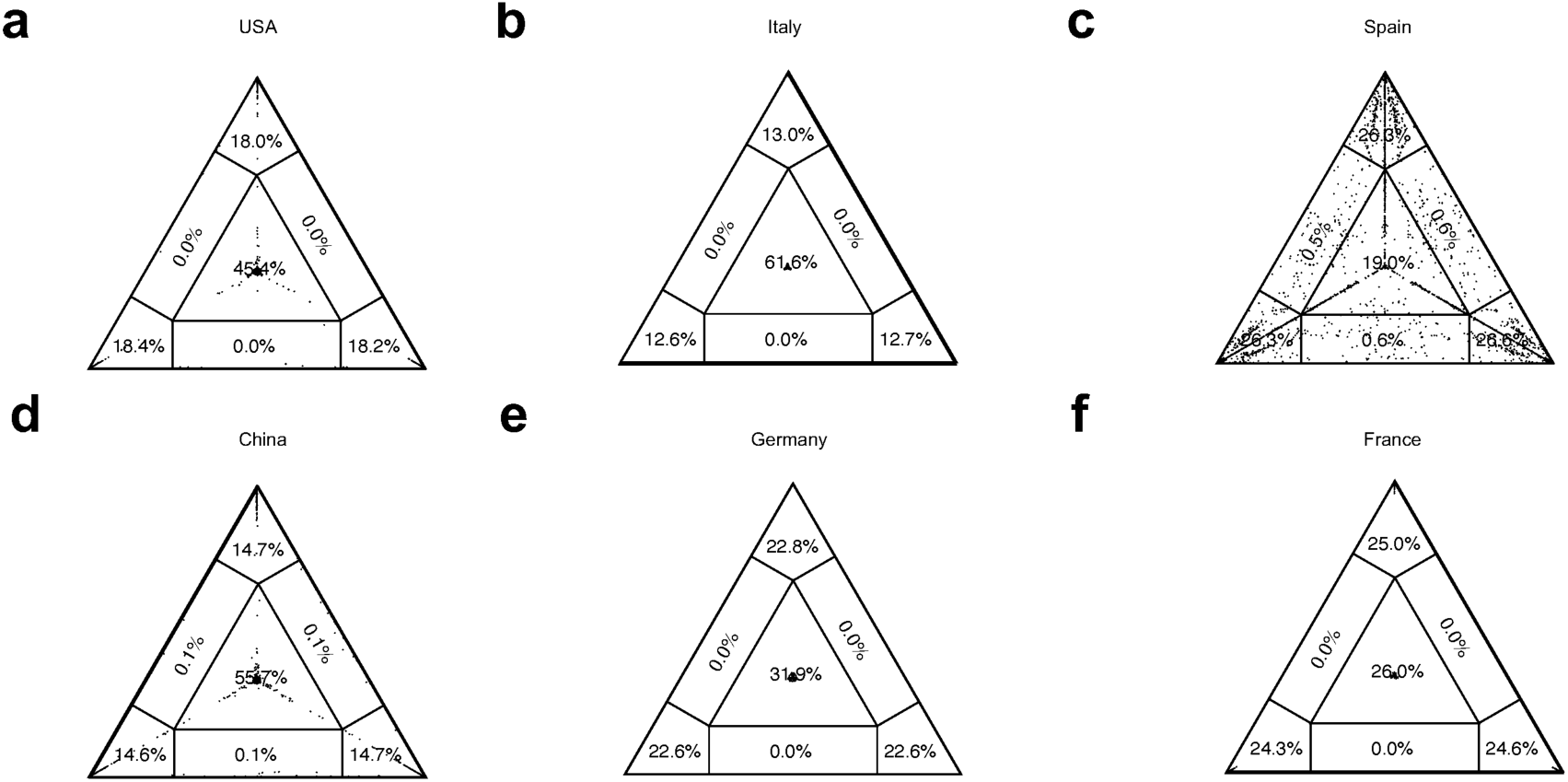
Phylogenetic signal for local SARS-CoV-2 datasets. Presence of phylogenetic signal was evaluated by likelihood mapping checking for alternative topologies (tips), unresolved quartets (center) and partly resolved quartets (edges) for genomes available on genome on March 30^th^ from (a) USA (612 genomes, 121,146 confirmed cases), (b) Italy (27 genomes, 97,689 confirmed cases), (c) Spain (40 genomes, 80,110 confirmed cases), (d) China (300 genomes, 82,122 confirmed cases), (e) Germany (27 genomes, 62,095 confirmed cases) and (f) France (119 genomes, 40,708 confirmed cases on March 29^th^). Presence of phylogenetic signal (<40% unresolved quartets in the center) was detected only for: Germany (27 genomes, 34 variant sites – 0.2% of total sites in the genome - 15 parsimony informative), with Düsseldorf and North Rhine Westphalia being the most contributing regions, respectively 12 and 11 genomes; Spain (40 genomes, 60 variant sites - 0.2% of total sites in the genome – 23 parsimony informative) with Madrid and Comunidad Valenciana being the most contributing regions, respectively 18 and 10 genomes; and France (119 genomes, 155 variant sites – 0.5% of total sites in the genome - 44 parsimony informative) with Auvergne-Rhône-Alpes, Hauts de France, and Bretagne being the most contributing regions, respectively 42, 30, and 13 genomes

**Figure S7.**
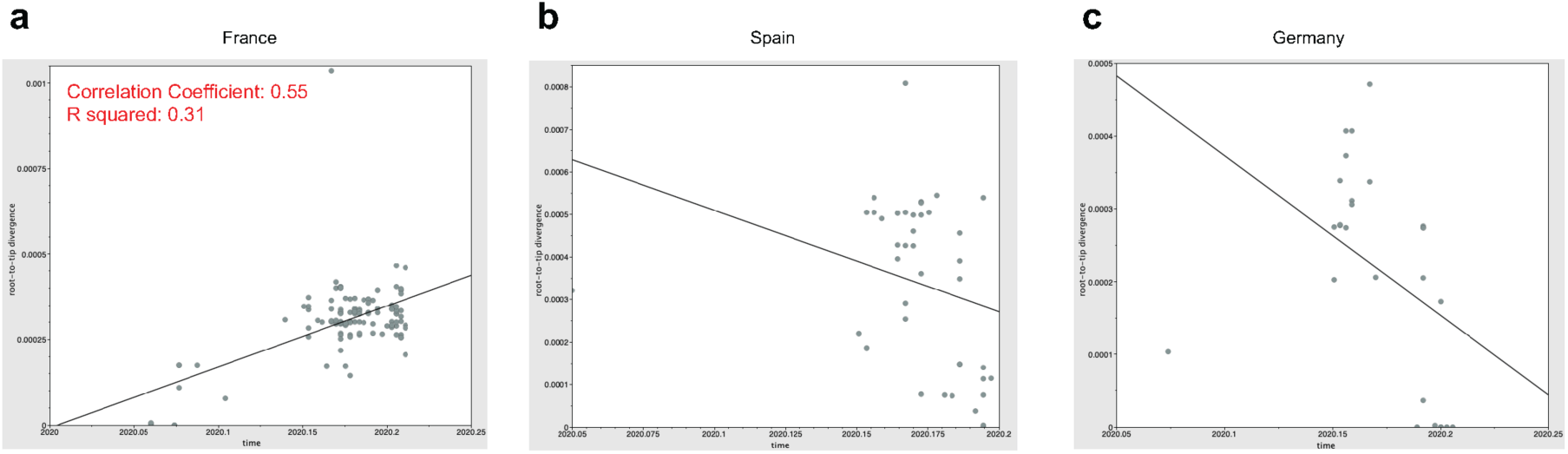
Regression analysis of temporal resolution of SARS-CoV-2 genomes datasets from France, Spain and Germany. The plots represent linear regression of root-to-tip genetic distance within the ML phylogeny against sampling time for each taxa. Temporal resolution was assessed using the slope of the regression, with positive slope indicating sufficient temporal signal for datasets collecet on (a) France, (b) Spain and (c) Germany. Correlation coefficient “r” are reported for each genomic fragment.

**Figure S8.**
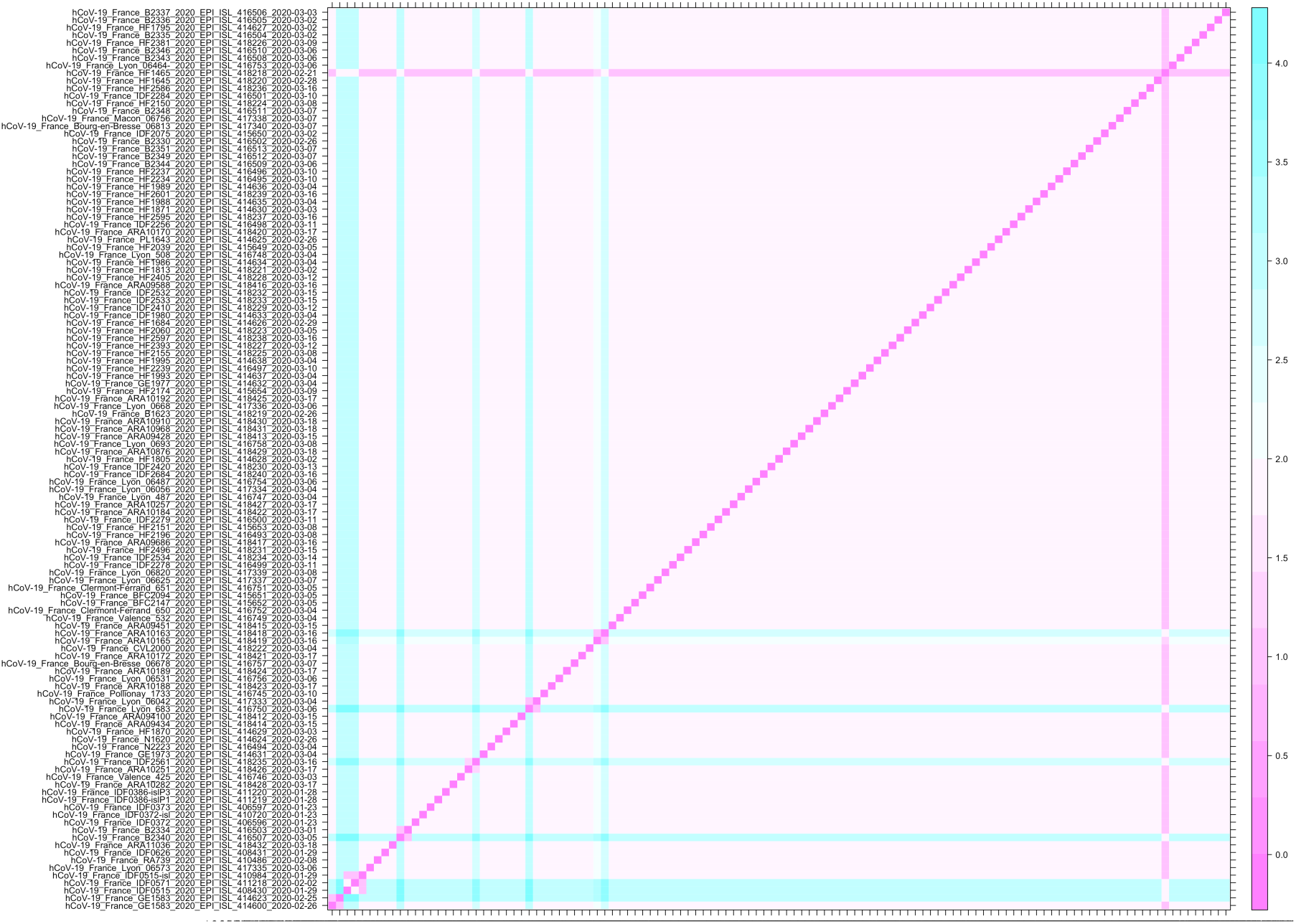
Transmission matrix of SARS-CoV-2 in France. Transmission matrix indicates for each pairs of individuals how many intermediates there are in the transmission chain are given for France (119 genomes as of March 30^th^ 2020, 40,708 confirmed cases as of March 29^th^ 2020).

**Table S1.** Acknowledgment table with full information of genome sequences. Downloaded from GISAID on March 10^th^ 2020 and provided as Excel file.

**Table S2.** Testing alternative topologies.

| Switched branches | Alternative Topology* | LogL** | $\Delta L^\#$ | SH $p$-value## |
| --- | --- | --- | --- | --- |
| Italy - Wales | 1 | -45443.2 | 0 | 0.24 |
| Germany - Brazil | 2 | -45451.5 | 8.3554 | 0.16 |
| *Portugal - Brazil* | 3 | -45443.2 | 0.0002 | 0.75 |
| *Germany - Portugal* | 4 | -45451.5 | 8.3197 | 0.16 |

\* Alternative topology obtained by switching branches in the ML tree inferred from SARS-CoV-2 full genome sequences. 1: Italy (EPI\_ISL\_412973) switched with Wales (EPI\_ISL\_413555); 2: Germany (EPI\_ISL\_406862) with Brazil (EPI\_ISL\_412964); 3: Portugal (EPI\_ISL\_413648) with Brazil (EPI\_ISL\_412964); 4: Germany (EPI\_ISL\_406862) with Portugal (EPI\_ISL\_413648). \*\* LogL: log likelihood estimated for each alternative topology. # $\Delta L$ : difference between LogL and the log likelihood of the original tree. ## SH: Shimodaira-Hasegawa test (22).

